# Pervasive false brain connectivity from electrophysiological signals

**DOI:** 10.1101/2021.01.28.428625

**Authors:** Roberto D. Pascual-Marqui, Peter Achermann, Pascal Faber, Toshihiko Kinoshita, Kieko Kochi, Keiichiro Nishida, Masafumi Yoshimura

## Abstract

Signals of brain electric neuronal activity, either invasively measured or non-invasively estimated, are commonly used for connectivity inference. One popular methodology assumes that the neural dynamics follow a multivariate autoregression, where the autoregressive coefficients represent the couplings among regions. If observation noise is present and ignored, as is common in practice, the estimated couplings are biased, affecting all forms of Granger-causality inference, both in time and in frequency domains. Significant nonsense coupling, i.e., nonsense connectivity, can appear when in reality there is none, since there is always observation noise in two possible forms: measurement noise, and activity from other brain regions due to volume conduction and low spatial resolution. This problem is critical, and is currently not being addressed, calling into question the validity of many Granger-causality reports in the literature. An estimation method that accounts for noise is based on an overdetermined system of high-order multivariate Yule-Walker equations, which give reduced variance estimators for the coupling coefficients of the unobserved signals. Simulation-based comparisons to other published methods are presented, demonstrating its adequate performance. In addition, simulation results are presented for a zero connectivity case with noisy observations, where the new method correctly reports no connectivity while classical analyses (as found in most software packages) report nonsense connectivity. For the sake of reproducible research, the supplementary material includes, in human readable format, all the time series data used here.

## 2. Introduction

Consider the multivariate autoregressive model (MAR) of order “*p*”, for the time series 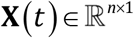:

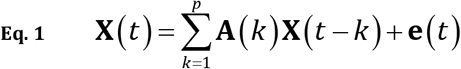

where 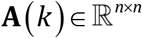 for *k* = 1..*p* are the autoregressive coefficients, and 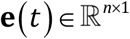 denotes the innovations with zero mean and innovations covariance matrix 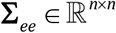.

Technical details on the multivariate autoregressive model can be found in Kilian and Lütkepohl (2017).

Assume that the actual measurements 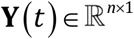 are contaminated with additive noise:

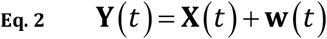

where **e**(*t*) and **w**(*t*) are independent, and 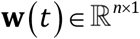 denotes the noise with zero mean and noise covariance matrix 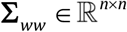.

The problem of interest is:

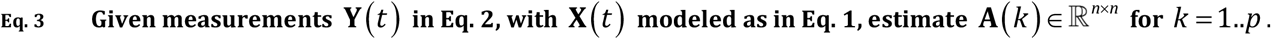

**X**(*t*) will be referred to as the unobserved signals, and **Y**(*t*) as the observed noisy signals or noisy data. Furthermore, **Σ**_*ee*_ is referred to as the innovations covariance, and **Σ**_*ww*_ as the noise covariance. Innovations are not to be confused with observation noise.

See e.g., Jamoos et al (2011), Mahmoudi and Karimi (2008), Qu et al (2011), and Rangarajan and Rao (2018) for the statement of this problem and several algorithmic solutions.

In formal terms, the null-hypotheses for testing Granger-causal-influence from region “*j*” to region “*i*”, as used in connectivity inference, correspond to the following two cases:

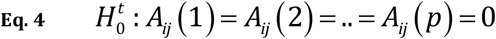

and:

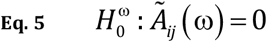

where *A_ij_*(*k*) corresponds to the ij-th element of **A**(*k*), and *Ã_ij_*(ω) corresponds to the complex valued discrete Fourier transform at frequency ω for the sequence *A_ij_*(*k*) for *k* = 1..*p*.

Evidence of Granger-causal connectivity corresponds to rejection of the null hypothesis Eq. 4 or Eq. 5.

In most brain connectivity studies based on the MAR model, it is implicitly assumed that there is no noise, and Eq. 1 is fitted directly to the measured or estimated signals of electric neuronal activity. See e.g., Astolfi et al (2007), Barnett and Seth (2014), Pascual-Marqui et al (2014), Seth et al (2015), Richter et al (2017), Coito et al (2019), Barnett et al (2020). In all cases, in the presence of observation noise, connectivity may arise (rejection of the null hypotheses) even when there is true independence.

In the next section, the method for estimating the connectivity parameters in an ideal, no noise case is presented first.

Next, the same equations are applied to observations in the presence of noise, which is an inappropriate procedure. This reveals the bias in the incorrectly estimated parameters.

Next, a simple example is worked out in detail, demonstrating how nonsense connectivity can appear in the case of signals of electric neuronal activity. This is due to the realistic fact that there is activity from other sources, which additively affect the measurements of interest, due to volume conduction and low spatial resolution.

Next, a solution to the problem in Eq. 3 is presented, denoted OHOYW: over determined high order Yule-Walker method for autoregressive coefficients estimation. See e.g. Friedlander (1983), and equation 2.138 in Najim (2008).

Finally, simulated data is used for the comparison of the present method with other published solutions, and for further illustrating the nonsense connectivity problem.

## 3. The classic multivariate autoregressive model

Let the expectation operator *E*[•] denote average over time samples. Then, post multiplying Eq. 1 by **X**^*T*^(*t* – *i*) and averaging gives:

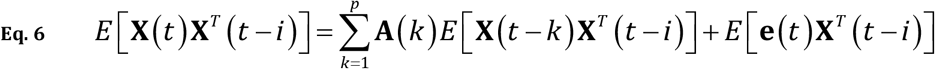

equivalent to:

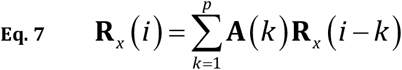

with:

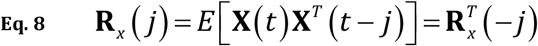

and:

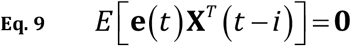

The Yule-Walker equations correspond to Eq. 7, for *i* = 1..*p*, and can be used for estimating the autoregressive coefficients **A**(*k*) for *k* = 1..*p*, given sample autocovariances **R**_*x*_(*j*) for *j* = 0..*p*. See e.g., equation 5.85 in Shumway and Stoffer (2017).

Eq. 9 follows from the causal independence of the present innovation **e**(*t*) with past values of **X**(*t* – *i*) for *i* > 0.

The Yule-Walker equations (Eq. 7), for *i* = 1..*p*, are:

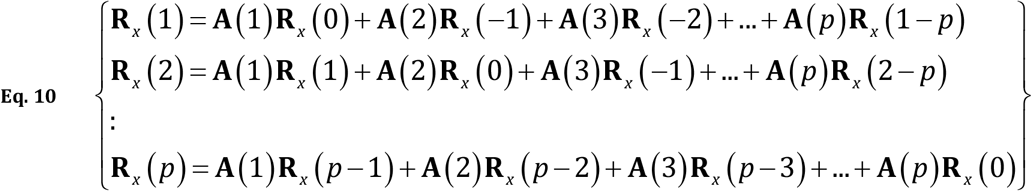

equivalent to:

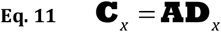

with:

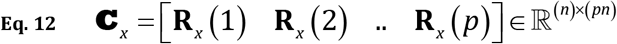

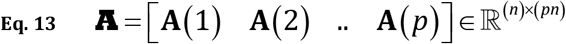

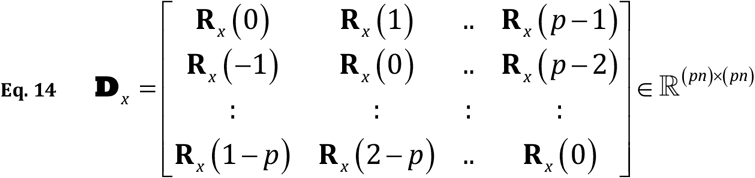

An estimator for the autoregressive coefficients is:

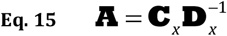

## 4. The biased estimators for the autoregressive parameters in the presence of noise

In this case the observations, i.e. the measurements, correspond to Eq. 2. The available data allow the computation of the autocovariances of **Y**(*t*), which would then be plugged into the Yule-Walker equations with autoregressive coefficients **B**(*k*):

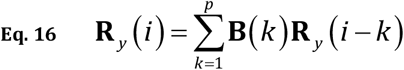

with:

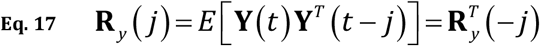

The relation between the autocovariances for the unobserved signals and the observed noisy signals are:

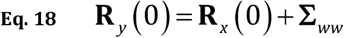

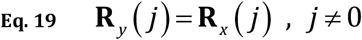

See e.g., equations 7 and 8 in Rangarajan and Rao (2018).

Eq. 16, Eq. 18, and Eq. 19 give (in analogy to Eq. 11, Eq. 12, Eq. 13, and Eq. 14):

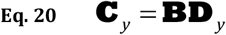

equivalent to:

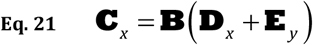

with:

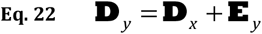

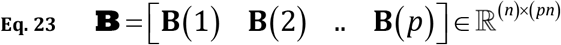

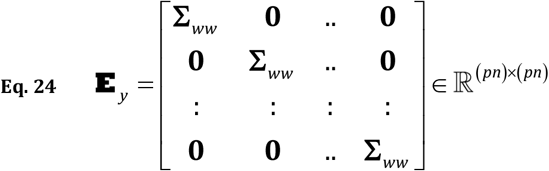

Note that in general, the autoregressive coefficients **B** obtained by solving Eq. 20 using noisy data, are different from the autoregressive coefficients **A** obtained by solving Eq. 11 using non-noisy data, i.e.:

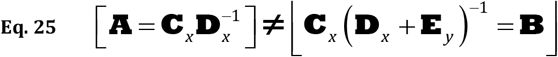

By definition, both coefficients **A** and **B** are identical in the noiseless case with Σ_*ww*_ = **0**, equivalent to **E**_*y*_ = **0**.

In practice, most publications are computing and making connectivity inference based on the biased autoregressive coefficients **B**.

The next section gives an explicit example of bias generating nonsense Granger causality from purely independent activity.

## 5. Nonsense connectivity

Consider the hypothetical case of two unobserved time series of cortical electric neuronal activity, that satisfy an autoregressive model of order *p*=1. It will be further assumed that the two signals are not interacting causally, i.e., the causal connectivity autoregressive coefficients in Eq. 27 and Eq. 28 are zero:

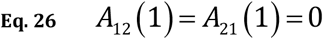

The model can be written as:

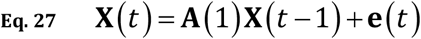

with 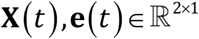, with autoregressive coefficients:

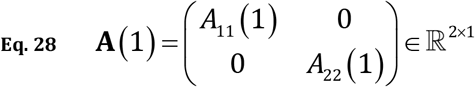

and with diagonal innovations covariance 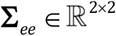.

Note that the autocovariance matrices **R**_*x*_(0) and **R**_*x*_(1) are, by construction, diagonal.

The measurements 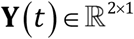 have added noise:

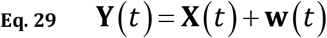

where it will be assumed that noise covariance is non-diagonal:

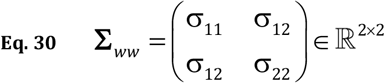

The autoregressive coefficients for the noisy data are obtained from Eq. 16, Eq. 18, and Eq. 19. which give:

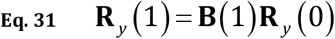

equivalent to:

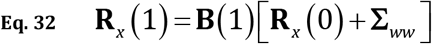

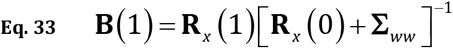

Define:

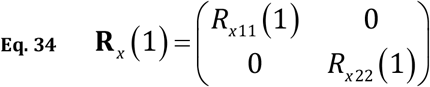

Then, from Eq. 30, Eq. 33, and Eq. 34, it follows that the estimated causal connectivity autoregressive coefficients are non zero:

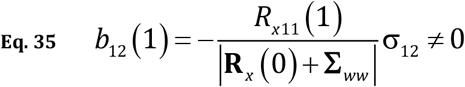

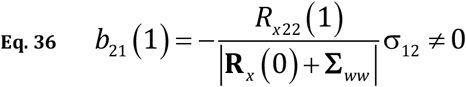

This demonstrates how nonsense, non-zero, false positive connectivity can occur: namely, by inappropriately fitting an autoregressive model to data contaminated by correlated observation noise.

Note that this situation can occur in real life for signals of electric neuronal activity, where the recordings are contaminated with activity from a third independent region due to volume conduction and low spatial resolution. In this case, the observation noise is very highly correlated since it is basically a common signal.

## 6. OHOYW: over determined high order Yule-Walker equations for noisy (and non-noisy) observations

The main difficulty in solving the problem in Eq. 3 is the unknown noise covariance Σ_*ww*_, for which there is no simple estimator. The number of publications on this multivariate problem is relatively scarce, but the diversity of algorithms is large, see e.g., Jamoos et al (2011), Mahmoudi and Karimi (2008), Qu et al (2011), and Rangarajan and Rao (2018). All these algorithms require an estimator for Σ_*ww*_, which is either updated iteratively or by means of the Kalman filter based on a state-space formulation of the problem.

The estimators presented here, for the autoregressive parameters of the unobserved signals, do not require an estimator for Σ_*ww*_.

The coupling coefficients, i.e. the autoregressive coefficients, for the unobserved signals are estimated directly from the autocovariances of the observed noisy data as shown next.

Consider the set of high order Yule-Walker equations (Eq. 7), for *i* = (*p* + 1)…*q*, with:

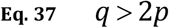

and:

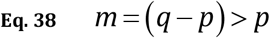

which gives *m>p* equations, for *p* coefficients:

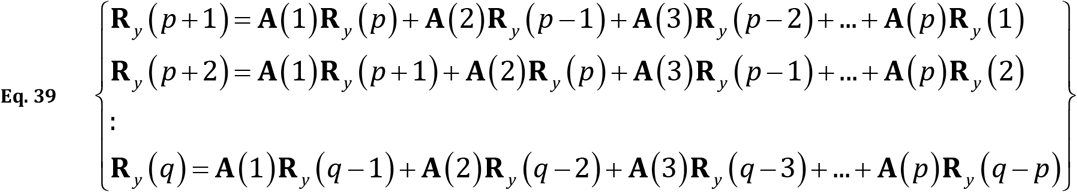

Use has been made here of the fact that the lagged autocovariances for the noisy observed data and for the unobserved signals are equal, as indicated in Eq. 19.

Eq. 39 is equivalent to:

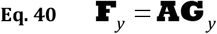

with:

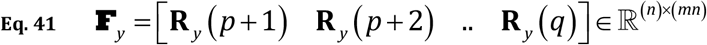

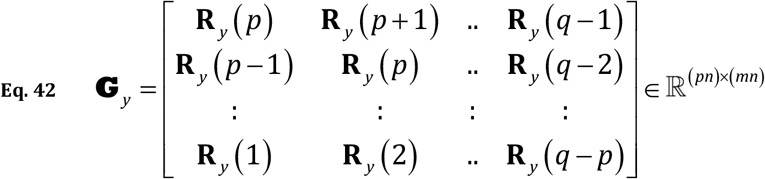

The least squares solution is:

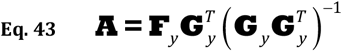

The number of equations used in this work is twice the number of autoregressive coefficients:

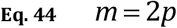

corresponding to:

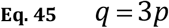

It has been argued, see e.g. Kay (1980), that the autoregressive coefficients estimated from the high order Yule-Walker equations have very high variance. This is certainly the case when the number of high order equations is limited to be equal to the number of coefficients. In our case, with the number of equations at least twice the number of parameters, the inverse matrix in Eq. 43 is much better conditioned than for the case of equal parameters and equations. This leads to estimators for the autoregressive coefficients with lower variance. See e.g. Friedlander (1983), and equation 2.138 in Najim (2008)

### Summary of the algorithm

Step#0: Referring to Eq. 1 and Eq. 2, and the problem defined in Eq. 3: given (possibly noisy) time series observations 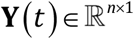, and the autoregressive order *p*.

Step#1: Compute:

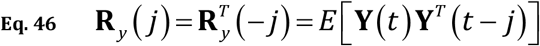

for *j* = (*p* + 1)… (3*p*). Plug Eq. 46 into Eq. 41 and Eq. 42, and solve Eq. 43. This gives the estimator **A** for the coupling (autoregressive) coefficients of the unobserved time series **X**(*t*) that appear in Eq. 1.

## 7. Comparison of the present method with other published solutions

Simulations from two publications are used here, namely from Mahmoudi and Karimi (2008), and from Diversi (2018), which study and compare the performance of several methods for solving the multivariate problem in Eq. 3. In all cases, noisy autoregressive models of order 2, for two time series, were used.

Two overall measures of quality were used in Mahmoudi and Karimi (2008), namely, the relative error (RE) and the normalized root mean squared error (nRMSE):

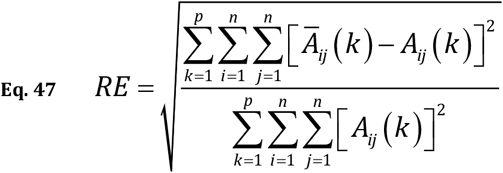

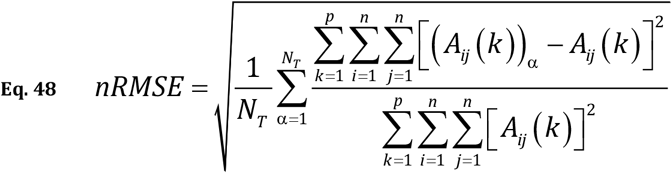

where *A_ij_*(*k*) is the true theoretical coupling coefficient at the k-th lag, from time series “*j*” to time series “*i*”, (*Â_ij_*(*k*))_*α*_ is its estimated value for the α-th trial, *Ā_ij_*(*k*) is the estimated average over all trials, and *N_T_* is the number of trials.

Although Diversi (2018) does not report relative error nor root mean squared error, the data presented in that publication is sufficient for the computation of relative errors.

Mahmoudi and Karimi (2008) perform three different simulations, consisting of 1000 trials, each with 4000 time samples, for:

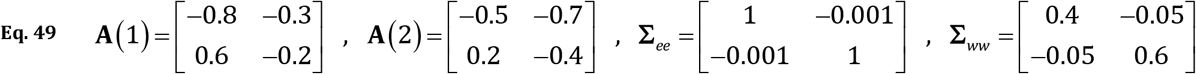

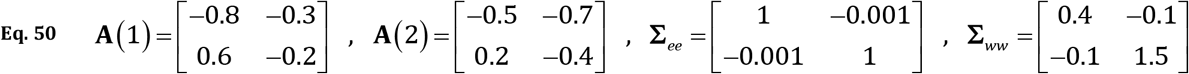

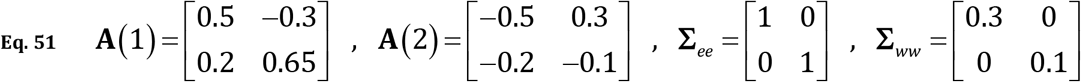

Diversi (2018) performs one simulation, consisting of 200 trials, each with 4000 time samples:

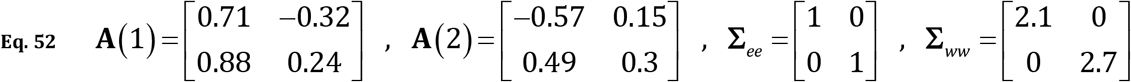

In all cases, innovations and noise have zero mean Gaussian distribution with the specified covariances.

Table 1 for the relative errors, and Table 2 for the root mean squared errors, show the comparative results for all methods studied in Mahmoudi and Karimi (2008) and for the method presented here.

**Table 1:** Relative errors (RE%) in the estimation of the autoregressive coefficients, for five methods, in three simulations (Eq. 49, Eq. 50, and Eq. 51). The simulations and the first four methods are described in Mahmoudi and Karimi (2008). The over determined high order Yule-Walker method for autoregressive coefficients (OHOYW) is the fifth method in the table.

| Method | Simulation#1 | Simulation#2 | Simulation#3 |
| --- | --- | --- | --- |
| LS | 24.0924 | 30.2876 | 16.9209 |
| MYW | 0.5058 | 0.833 | 10.7929 |
| ILSV | 1.1981 | 3.0951 | 0.9639 |
| HasanEtAl | - | - | 1.1023 |
| OHOYW | 0.78015 | 1.0744 | 1.5259 |

LS: least squares, MYW: modified Yule-Walker, ILSV: improved least-squares for vector processes, all referenced in Mahmoudi and Karimi (2008). HasanEtAl: Hasan et al (2003). OHOYW: over determined high order Yule-Walker method for autoregressive coefficients (method presented here).

**Table 2:** Normalized root mean squared error (nRMSE%) in the estimation of the autoregressive coefficients, for five methods, in three simulations (Eq. 49, Eq. 50, and Eq. 51). The simulations and the first four methods are described in Mahmoudi and Karimi (2008). The over determined high order Yule-Walker method for autoregressive coefficients (OHOYW) is the fifth method in the table.

| Method | Simulation#1 | Simulation#2 | Simulation#3 |
| --- | --- | --- | --- |
| LS | 24.2461 | 30.4298 | 17.3643 |
| MYW | 13.7287 | 17.4084 | 327.4975 |
| ILSV | 6.8094 | 11.1532 | 10.5154 |
| HasanEtAl | - | - | 11.6721 |
| OHOYW | 12.829 | 15.346 | 43.722 |

LS: least squares, MYW: modified Yule-Walker, ILSV: improved least-squares for vector processes, all referenced in Mahmoudi and Karimi (2008). HasanEtAl: Hasan et al (2003). OHOYW: over determined high order Yule-Walker method for autoregressive coefficients (method presented here).

Table 3 shows the comparison of relative errors for all methods studied in Diversi (2018) and for the method presented here.

**Table 3:**
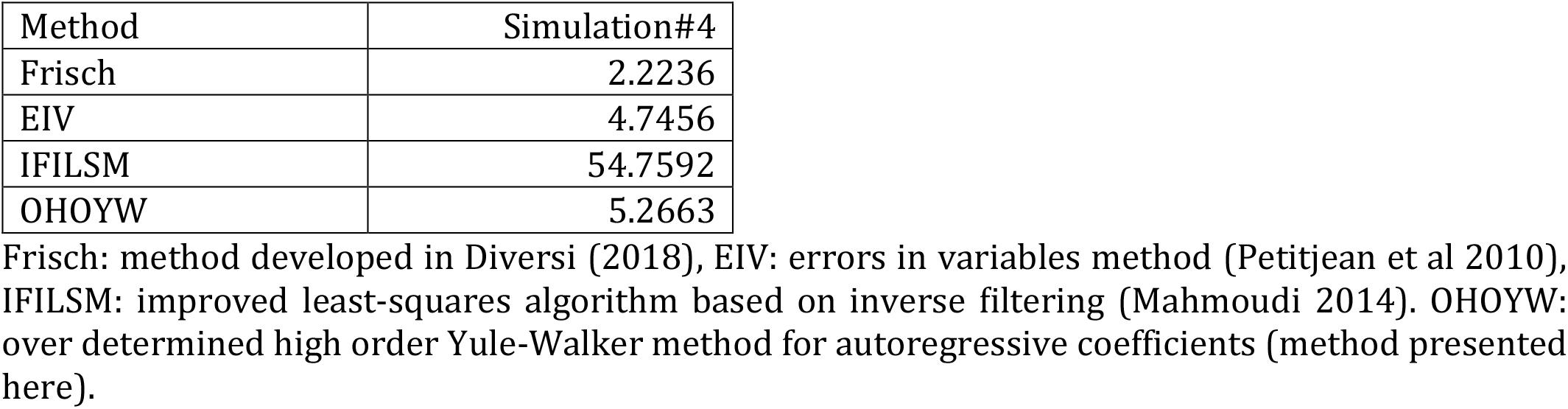
Relative errors (RE%) in the estimation of the autoregressive coefficients, for four methods, in one simulation (Eq. 52). The simulations and the first three methods are described in Diversi (2018). The over determined high order Yule-Walker method for autoregressive coefficients (OHOYW) is the fourth method in the table.

## 8. Simulations related to cases of true independence (no-connectivity): from nonsense connections with classical algorithms to correct inference with the new algorithm

Consider two unobserved independent time series modeled as:

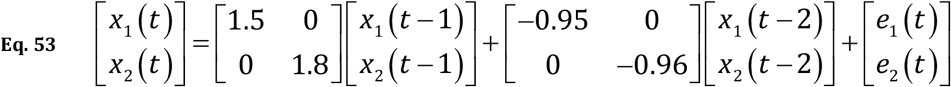

with noisy observations:

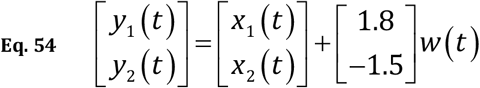

with **Σ**_*ee*_ = **I** and singular noise covariance:

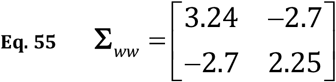

Note, from Eq. 54 and Eq. 55, that the noise consists of one single common source, affecting the measurements of both time series.

Simulations consisted of 100 trials, each with 256 time samples. In all cases, innovations and noise have zero mean Gaussian distribution with the specified covariances. Table 4 shows statistics of the estimated model when using OHOYW (over determined high order Yule-Walker method).

**Table 4:** Summary statistics of the estimated model when using OHOYW, for the simulation with parameters defined in Eq. 53, Eq. 54, and Eq. 55, based on 100 trials, each with 256 time samples.

|  | AR(1) |  |  | AR(2) |  |
| --- | --- | --- | --- | --- | --- |
| True AR coeffs | 1.5000 | 0.0000 |  | -0.9500 | 0.0000 |
|  | 0.0000 | 1.8000 |  | 0.0000 | -0.9600 |
| Means | 1.4920 | -0.0048 |  | -0.9441 | 0.0042 |
|  | -0.0020 | 1.7973 |  | -0.0009 | -0.9559 |
| StdDevs | 0.0416 | 0.0270 |  | 0.0355 | 0.0245 |
|  | 0.0324 | 0.0272 |  | 0.0377 | 0.0273 |
| RE(%) | 0.4801 |  |  |  |  |
| nRMSE(%) | 3.3968 |  |  |  |  |

True AR coeffs: true autoregressive coefficients, as in Eq. 53; Means: mean values over 100 trials; StdDev: standard deviations over 100 trials; RE(%): relative errors; nRMSE(%): normalized root mean squared error.

Table 5 shows statistics of the estimated model when using classical least squares estimation, which does not consider possible noise in the observations.

**Table 5:** Summary statistics of the estimated model when using classical least squares estimation, which does not consider possible noise in the observations, for the simulation with parameters defined in Eq. 53, Eq. 54, and Eq. 55, based on 100 trials, each with 256 time samples.

|  | AR(1) |  |  | AR(2) |  |
| --- | --- | --- | --- | --- | --- |
| True AR coeffs | 1.5000 | 0.0000 |  | -0.9500 | 0.0000 |
|  | 0.0000 | 1.8000 |  | 0.0000 | -0.9600 |
| Means | 1.1081 | 0.2840 |  | -0.6008 | -0.2737 |
|  | 0.3686 | 1.5270 |  | -0.3171 | -0.7007 |
| StdDevs | 0.1311 | 0.1097 |  | 0.1232 | 0.1073 |
|  | 0.1166 | 0.1128 |  | 0.0996 | 0.1105 |
| RE(%) | 33.2630 |  |  |  |  |
| nRMSE(%) | 35.3430 |  |  |  |  |

True AR coeffs: true autoregressive coefficients, as in Eq. 53; Means: mean values over 100 trials; StdDev: standard deviations over 100 trials; RE(%): relative errors; nRMSE(%): normalized root mean squared error.

## 9. Discussion

Table 1 and Table 3 show the relative errors in the estimated autoregressive coefficients for a wide range of published algorithms that deal with noisy time series observations. There are four simulations from the literature, and in all cases the simple, non-iterative method of overdetermined high order Yule-Walker equations (OHOYW: Friedlander (1983), and Najim (2008)) ranks very high in terms of performance.

Table 2 shows the root mean squared error as a comparative performance measure, and in this case, the OHOYW method is second best in two out of five simulations. In the third simulation OHOYW comes out in fourth place (out of five).

In summary, given the favorable comparative performance of the OHOYW estimators, and given the simplicity of the method, its use is recommended when estimating multivariate autoregressive coefficients in electrophysiology.

The simulation results in Table 4 and Table 5 demonstrate one important point: The straightforward use of multivariate autoregressive modeling assuming no noise can produce nonsense brain connectivity results, which is the case of almost all previous published results. This is apparent from relatively low standard deviations of the coupling coefficients from Table 5, which are at least two times smaller than the coupling value, thus confirming non-zero significant nonsense connectivity.

Also noteworthy from Table 4 and Table 5 are the performance measures, which show root squared mean error and relative error roughly 10 to 100 times worse for the least squares method.

This problem is solved with proper modeling and estimation, using the OHOYW equations.

The results shown in this work justifies the use of one of the two following procedures for connectivity inference: Procedure#1 (confirmatory): Estimate the multivariate autoregressive coefficients using the OHOYW equations.

Procedure#2 (exploratory): Perform two separate estimations of the multivariate autoregressive coefficients. One based on the classic least squares method that does not correct for noise, and the other using the OHOYW method. If the results are significantly different, use the OHOYW estimators.

Actually, with respect to “Procedure#2”, a test for equal autoregressive coefficients between the two methods (least squares and OHOYW) can be carried if the number of trials is relatively large.

There are some practical instances when it can be safely assumed that the noise is negligible. This might be the case of ECoG recordings from electrode arrays, when the analysis is based on local bipolar signals, and not on the raw signals, as demonstrated in Trongnetrpunya et al (2016). Local bipolars will be very little affected by observation noise originating from activity outside the vicinity of the bipolars, which is, in practice, the main cause of nonsense connectivity.

## 10. Limitation

Most of the literature on this problem of noisy measurements assumes that the observation noise is white, in the sense that it is not serially correlated. In electrophysiology, due to volume conduction and low spatial resolution, this assumption is violated, since electric neuronal activity from other regions is not white in general. The quantitative effect of such a violation needs further study.

## References

L. Astolfi, F. Cincotti, D. Mattia, M.G. Marciani, L.A. Baccala, F. de Vico Fallani, S. Salinari, M. Ursino, M. Zavaglia, L. Ding. Comparison of different cortical connectivity estimators for high-resolution EEG recordings. Hum. Brain Mapp., 28 (2) (2007), pp. 143–157

Barnett L, Muthukumaraswamy SD, Carhart-Harris RL, Seth AK. Decreased directed functional connectivity in the psychedelic state. NeuroImage. 2020 Apr 1; 209:116462.

Barnett L, Seth AK. The MVGC multivariate Granger causality toolbox: a new approach to Granger-causal inference. Journal of neuroscience methods. 2014 Feb 15; 223: 50–68.

Coito A, Michel CM, Vulliemoz S, Plomp G. Directed functional connections underlying spontaneous brain activity. Human brain mapping. 2019 Feb 15;40(3):879–88.

R. Diversi: Identification of Multichannel AR Models with Additive Noise: a Frisch Scheme Approach. 2018 26th European Signal Processing Conference (EUSIPCO), Rome, Italy, 2018, pp. 1252–1256, doi: 10.23919/EUSIPCO.2018.8553415.

Friedlander B. Instrumental variable methods for ARMA spectral estimation. IEEE Transactions on Acoustics, Speech, and Signal Processing. 1983 Apr;31(2):404–15.

Hasan MK, Hossain MJ, Haque MA. Parameter estimation of multichannel autoregressive processes in noise. Signal processing. 2003 Mar 1;83(3):603–10.

Jameson A. Solution of the equation AX+XB=C by inversion of an M*M or N*N matrix. SIAM Journal on Applied Mathematics. 1968; 16(5):1020–3.

Jamoos A, Grivel E, Shakarneh N, Abdel-Nour H. Dual optimal filters for parameter estimation of a multivariate autoregressive process from noisy observations. IET Signal Processing. 2011; 5(5):471–9.

Kay S. Noise compensation for autoregressive spectral estimates. IEEE Transactions on Acoustics, Speech, and Signal Processing. 1980; 28(3):292–303.

Kilian L, Lütkepohl H. Structural vector autoregressive analysis. Cambridge University Press; 2017.

Mahmoudi A. Adaptive algorithm for multichannel autoregressive estimation in spatially correlated noise. Journal of Stochastics. 2014 Jun 19;2014.

Mahmoudi A, Karimi M. Estimation of the parameters of multichannel autoregressive signals from noisy observations. Signal Processing. 2008; 88(11):2777–83.

Najim M. Modeling, estimation and optimal filtering in signal processing. J. Wiley & Sons; London, 2008.

Pascual-Marqui RD, Biscay RJ, Bosch-Bayard J, Lehmann D, Kochi K, Kinoshita T, Yamada N, Sadato N. Assessing direct paths of intracortical causal information flow of oscillatory activity with the isolated effective coherence (iCoh). Frontiers in human neuroscience. 2014 Jun 20; 8:448.

Petitjean J, Grivel E, Bobillet W, Roussilhe P. Multichannel AR parameter estimation from noisy observations as an errors-in-variables issue. Signal, Image and Video Processing. 2010 Jun;4(2):209–20.

Qu X, Zhou J, Luo Y. A new noise-compensated estimation scheme for multichannel autoregressive signals from noisy observations. The Journal of Supercomputing. 2011; 58(1):34–49.

Rangarajan P, Rao RP. Estimation of vector autoregressive parameters and granger causality from noisy multichannel data. IEEE Transactions on Biomedical Engineering. 2018; 66(8):2231–40.

Richter CG, Thompson WH, Bosman CA, Fries P. Top-down beta enhances bottom-up gamma. Journal of Neuroscience. 2017 Jul 12;37(28):6698–711.

Seth AK, Barrett AB, Barnett L. Granger causality analysis in neuroscience and neuroimaging. Journal of Neuroscience. 2015 Feb 25;35(8):3293–7.

Shumway RH, Stoffer DS. Time series analysis and its applications: with R examples. Fourth Edition. Springer. 2017.

Trongnetrpunya A, Nandi B, Kang D, Kocsis B, Schroeder CE, Ding M. Assessing granger causality in electrophysiological data: removing the adverse effects of common signals via bipolar derivations. Frontiers in systems neuroscience. 2016 Jan 20;9:189.

