## Supplementary material for "Pervasive false brain connectivity from electrophysiological signals": diversi2018params.txt

2			
2			

1 0
0 1

+0.71  -0.32
+0.88 +0.24

 -0.57 +0.15
+0.49 +0.30

 2.1 0
 0   2.7


R. Diversi: Identification of Multichannel AR Models with Additive Noise: a Frisch Scheme Approach. 2018 26th European Signal Processing Conference (EUSIPCO), Rome, Italy, 2018, pp. 1252-1256, doi: 10.23919/EUSIPCO.2018.8553415.
Note that the ARcoeffs are defined as negative in his convetion!
nt=4000			
nepochs=200			
