## Supplementary material for "Pervasive false brain connectivity from electrophysiological signals": SimulationsTable4Table5-params.txt

2			
2			

1 0
0 1

 1.5 0
 0 1.8
 
 -0.95 0
 0 -0.96
 
3.24 -2.7
-2.7 2.25

nepochs=100
number of time samples for each epoch=256

simulation data as in table 4 and table 5 from:
Pascual-Marqui et al. Pervasive false brain connectivity from electrophysiological signals. arxiv.org birxiv.org 2021-01
